# *eProbe*: a capture probe design toolkit for genetic diversity reconstructions from ancient environmental DNA

**DOI:** 10.1101/2024.09.02.610737

**Authors:** Zihao Huang, Zhengquan Gu, Yuanyang Cai, Ruairidh Macleod, Zhe Xue, Haoran Dong, Søren Overballe-Petersen, Shanlin Liu, Yu Gao, Hao Li, Sha Tang, Xianmin Diao, Morten Egevang Joergensen, Christoph Dockter, Lasse Vinner, Eske Willerslev, Fahu Chen, Hongru Wang, Yucheng Wang

## Abstract

Ancient environmental DNA (aeDNA) is now commonly used in paleoecology and evolutionary ecology, yet due to difficulties in gaining sufficient genome coverage on individual species from metagenome data, its genetic perspectives remain largely uninvestigated. Hybridization capture has proven as an effective approach for enriching the DNA of target species, thus increasing the genome coverage of sequencing data and enabling population and evolutionary genetics analysis. However, to date there is no tool available for designing capture probe sets tailored for aeDNA based population genetics. Here we present *eProbe*, an efficient, flexible and easy-to-use program toolkit that provides a complete workflow for capture probe design, assessment and validation. By benchmarking a probe set for foxtail millet, an annual grass, made by the *eProbe* workflow, we demonstrate a remarkable increase of capturing efficiency, with the target taxa recovery rate improved by 577-fold, and the genome coverage achieved by soil capture-sequencing data even higher than data directly shotgun sequenced from the plant tissues. Probes that underwent our filtering panels show notably higher efficiency. The capture sequencing data enabled accurate population and evolutionary genetic analysis, by effectively inferring the fine-scale genetic structures and patterns, as well as the genotypes on functional genes.

## 1. Introduction

Ancient DNA (aDNA, the analysis of DNA molecules preserved in remains of ancient organisms) has extensively advanced insights into many evolutionary and ecological questions, including the demographic dynamic of human populations [1,2,3,4], the adaptation and domestication of animals [5,6,7,8], and the coevolutionary trajectories of pathogens [9,10]. However, aDNA research has thus far relied predominantly on the isolation of DNA from well-preserved fossilised specimens, typically bone, hair, and teeth. Sampling is therefore limited to organisms with extensive hard-tissue (namely vertebrates, and occasionally trees or corals), and has so far centred on humans and domesticated animals, excluding the vast majority of species diversity like plants, small and soft body animals, protists, and microbes. Furthermore, excavated fossils frequently present a highly discontinuous temporal and spatial record, while genetic data representing significant chronological continuity can provide drastically improved insights into evolutionary processes [11,12]. Moreover, most fossil materials are unique and therefore archaeologically and culturally valuable. Ethical concerns have arisen in recent years for the invasive and destructive sampling of fossils for DNA extractions [13], as these unique materials hold the potential to yield greater insights with more advanced techniques in the future.

Along with the rapid technical advances in the past decade, ancient environmental DNA has emerged as a novel approach to study aDNA [14,15]. aeDNA refers to the ancient DNA molecules preserved in different types of environmental substrates, typically buried sediments such as lake and marine sediments [16,17,18], cave and archaeological deposits [19,20,21], and permafrost [22,23,24], but also other materials such as ice core and speleothems [25,26]. aeDNA uses ubiquitous environmental materials that are available in large amounts in both human formed depositions and natural depositional profiles across cryospheric, temperate, and tropical landscapes under different terrestrial, lacustrine, and marine sedimentation settings [15,27]. It usually forms continuous depositions offering successive records in high temporal resolutions (e.g., lake sediments can easily resolve the successional patterns in decades or even higher temporal resolutions [28,29,30]. Furthermore, it has been proven that aDNA can be preserved for a longer time in sediments compared to in bones in the same site [19,31], partially due to DNA being able to bind with the mineral substances in soil to form stable structures [32]. aeDNA therefore holds a great potential allowing for investigating into the deep past of evolutionary processes [24]. Moreover, the recovered sequencing data, as a pool of mixed DNA, covers the genetic information of taxa from all trophic levels [16,17,24], thus extending the scope of aDNA study to all organismal groups. These features combined situate aeDNA as a unique resource for extensively studying the ancient genomics and evolutionary genetics universally for organisms.

Conversely, aeDNA has so far been predominantly employed for reconstructing taxonomic composition [22,24] and tracking biotic successions in different ecosystems [16,17,23,28,29,30]. Due to the inherent complexity of aeDNA sources and substantial background noise caused by the overwhelming presence of microbial DNA in sediments [15,33], acquiring sufficient informative DNA sequences for the genomes of specific organisms is extremely challenging. Consequently, the potential genetic and genomic insights aeDNA preserves are rarely investigated, limiting its application on phylogenetic, evolutionary, and population genetic analysis. One effective approach to tackle this problem, and hence to improve the specificity and efficiency of aeDNA sequencing, is targeted sequencing, whereby specific DNA sequences are selectively enriched through hybridization capture [34]. Despite the capture approach having been extensively applied in fossil-based aDNA studies [35,36], facilitating most attempts in resolving the genetic information at population level using aeDNA [37,38,39], no tool currently exists for designing and accessing the capture probe sets tailored for population genetic research, which forms the very first step to initialise a capture experiment for aeDNA study. The main focus of published capture aeDNA studies has not been placed on the design of probe sets [18,37,38,39,40,41]. Research efforts in this line of work have been centred on experimentally improving [42] or empirically examining the capture efficiency [39,41], capture baits synthesis and generation [43,44], and designing probes for capturing the biodiversity of target loci with broad taxonomic applicability [45,46,47,48].

To fix this gap, here we introduce *eProbe*, a Python package tailored for designing probe sets for target genome capture sequencing, specifically accounting for the features of aeDNA. Facilitated by the *eProbe* package, we present a complete workflow that enables effectively generating, filtering and selecting probe sequences, as well as assessing and validating the probe set from both experimental and analytical perspectives. We further benchmarked the workflow by designing and synthesising a capture bait set targeting the population history and functional gene variation of foxtail millet (*Setaria italica*) and wild foxtail millet (*S. viridis*). It showed that the probe set remarkably improved the sequencing efficiency for the target taxa by an average of 455-fold, and the capture-sequenced data enabled to elucidate fine-scale population genetic structures.

## 2. The *eProbe* package and workflow

The overall idea of the *eProbe* workflow aims at capturing informative single nucleotide polymorphisms (SNPs) across the genome of targeted species. eProbe designs probe sets for capturing the genetic diversity of a single species or a series of closely related taxa. However, it’s not taxonomically sensitive, and can be applied for designing probe sets for any genomic regions of interest.

*eProbe* generates two separate probe set panels based on their research objectives and the genomic regions of interest. The main panel is for inferring the population genetic patterns (thus the POPGEN panel) that will go through the filterings for selecting the highest quality and best representative probes. However, many key functional genes of general interests are conserved among species. The genome regions of the target taxa covering these genes therefore share high homology with other taxa, resulting in most probes designed for these genes will subsequently be filtered out for low specificity. The second panel (referred as the FUNCGEN panel) is therefore designed to cover these functional genes, or any genome regions that will not be subjected to the filtering steps. The two panels are ultimately merged together as the final probe set. Here we describe the full application processes along with the complete workflow (Fig. 1).

**Fig. 1.**
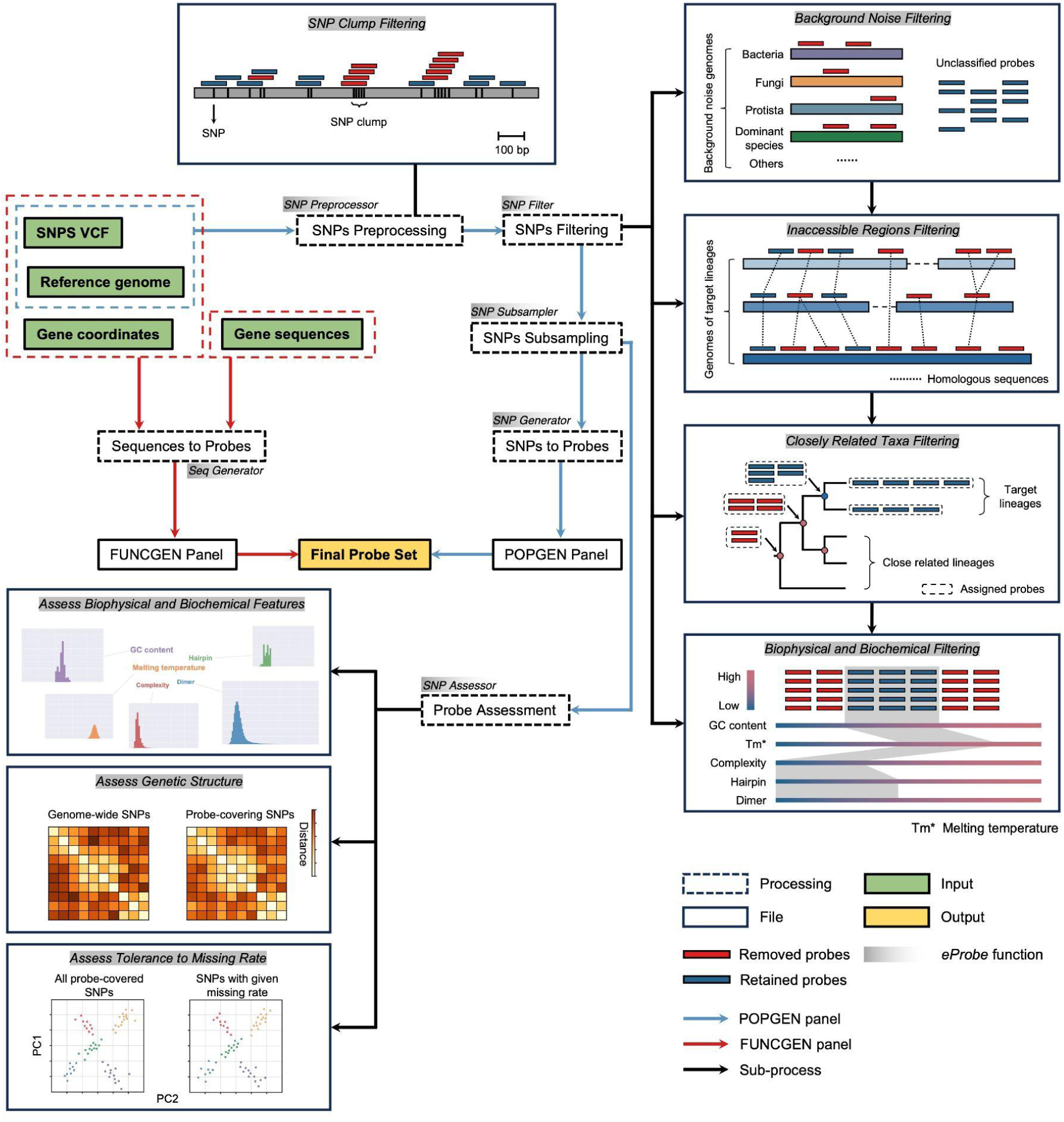
The *eProbe* workflow. Depending on the desired probe set panels, different input files are required. The dash blue rectangle encircles the input files for the POPGEN panel, while the red blue rectangles encircle different input formats for the FUNCGEN panel.

### 2.1 Input data

The POPGEN panel requires two input files: a reference genome in FASTA format and a collection of single nucleotide polymorphisms (SNPs, genetic variants) in VCF format. We only kept bi-allelic SNPs in the VCF input for the following analyses as a common practice in most genetic structure and population genetics inferences [49,50,51,52,53]. Samples (or individuals) included in the VCF file should represent the genetic diversity of the target species across their known spatial distribution, and ideally also cover candidate ancestral-state variants through sampling of multiple diverged descendants. We also present a pipeline for generating the input VCF file from the user’s own genome sequencing data of relevant samples or individuals for the target species (see [*eProbe* GitHub repository](https://github.com/YCWangLab/eProbes)).

The FUNCGEN panel requires gene sequences that can be supplied in various input formats, such as the sequences (in FASTA format), the genomic coordinates (in BED format), or the variants in genic regions (in VCF format), which is used for obtaining different haplotypes of functional genes. The two pipelines can be executed independently, and users can supply files for any of the two pipelines to initialise the process.

### 2.2 Probes for the POPGEN panel

#### 2.2.1 SNPs preprocessing

SNP clusters (*i.e.*, multiple SNPs that are closely clustered together on genomes) are commonly observed across species (see Supplementary Information). Placing probes on SNP clusters increases divergence between probes and target sequences, thereby introducing uncenterties and bias towards alleles more similar to those in the template genome. Avoiding SNP clusters is particularly important for aeDNA probes design since both deamination damage and mutations caused by evolutionary divergences of aeDNA further introduce mismatches against the modern reference genomes, which increases the bias. *eProbe* therefore supplies a function (*SNP Preprocessor*) to first filter out SNP clusters in the supplied VCF files. This function also allows to retain or exclude SNPs from specific genomic regions.

#### 2.2.2 SNPs filterings

There are normally millions of informative loci (specific sites on a chromosome) across a given genome of higher plants and animals, while the majority are not suitable as anchor sites for capture probes due to severe interference from abundant non-target sequences, either from the environment or the genome itself, poor performance during hybridization capture, or the captured sequences being unusable for analysis. *eProbe* supplies a series of functions to detect and filter loci, by removing SNPs with flanking sequences found in other taxa across a comprehensive reference genome database and in close-related taxa, in inaccessible genomic regions (see Supplementary Information), and importantly in genomic regions with low quality and/or outlier values for their biophysical and biochemical features (see Supplementary Information). All filterings are optional, and different filtering functions can be combined in different orders for flexible filterings.

##### 2.2.2.1 Filtering background noises

As a mixed pool of DNA originated from thousands of organisms, the endogenous rate of the target taxa directly yielded from aeDNA are usually extremely low [20]. The most significant noise that limits the efficiency of aeDNA capture comes from the background DNA sequences of other taxa, including microbes as the majority and also non-target eukaryotes of high abundance. *eProbe*’s *SNP Filter* function supplies an aggressive *k*-mer based approach (using Kraken 2 [54]) to fast and sensitively screen all possible probe candidates around each SNP against large genome reference databases (such as GTDB, NCBI-nt, NCBI-refseq, or any custom-modified genome databases), and remove SNPs flanked by sequences that show high similarity score with any non-target taxa.

##### 2.2.2.2 Filtering inaccessible genome regions

Structural variations (SVs, such as insertions, deletions, and duplications) are commonly found among individuals and populations, resulting in an inaccessibility of genomic regions for certain lineages of given species [55,56]. Further, DNA sequences from repetitive regions in a broad range of organisms [57] can hardly be aligned to a reference genome rendering it useless for subsequent analysis. These genomic regions are collectively referred to as inaccessible regions. Using a series of flexible parameters, *eProbe* employs an alignment-based approach (included in the function *SNP Filter*) to detect and remove SNPs in these problematic genomic regions, by requiring the probe candidates to be able to align to the selected and suitable genomic regions (see Supplementary Information).

##### 2.2.2.3 Filtering against closely related taxa

Phylogenetically closely-related taxa usually share a large amount of identical sequences (*e.g.*, 98.8% of human and chimpanzee genomes are identical [58]. Distinguishing homologous sequences among these taxa is important but challenging, particularly in the archaeological context where closely-related taxa (*e.g.*, domesticated crop *versus* wild grass) usually coexist. *eProbe* provides an alignment-based approach (included in the function *SNP Filter*) that enables the user to retain SNPs in regions exclusively assignable to the target species through comparison against reference genomes of closely related taxa.

##### 2.2.2.4 Filtering on probes’ biophysical and biochemical features

The biophysical and biochemical features of the probe sets determine the performance of hybridization reactions and thus the efficiency of the final synthesised capture baits. In this step, we consider four different features for filtering the probe candidates: 1) the GC content and melting temperature (Tm) of probe sequences that would affect the specificity and stability of hybridization combinations of the synthesised capture baits (ideally Tm for a probe set should be narrowly distributed so that a uniform annealing temperature can be adopted for hybridization reaction [59], 2) complexity of the probe sequences (*e.g.*, rate of homopolymer, dinucleotide and trinucleotide repeats) that affects the rate of non-specific hybridization combinations [60], 3) propensity for forming hairpin structures for each probe *(i.e.*, the probe itself not containing reverse complementary sequences), and 4) predicted likelihood of forming probe dimers between each pair across all probes *(i.e.*, avoiding reverse complementary sequences among probes). *eProbe* supplies functions (included in function *SNP Filter*) to first quickly assign scores on each of the four filtering features for all possible probe candidates, and then based on the scoring matrix to filter out probes with low quality and/or outlier values based on user-defined thresholds for each feature.

#### 2.2.3 Selective subsampling and probe set generating

After the above-discussed filterings, SNPs can be further subsampled into any custom size (function *SNP Subsampler*) to match the specific quantity requirements of the probe set. To ensure an even coverage on the reference genome, *eProbe* employs a sliding window approach to extract SNPs that have passed all filterings in each window distributed throughout the entire reference genome. Within each window, *eProbe* allows: 1) random selection of a SNP, 2) optimization of SNP selection through revisiting the scoring matrix of probes’ biophysical and biochemical features, and 3) prioritisation of SNPs in regions of interest (details in Supplementary Information).

The final probe set for the POPGEN panel can be generated using the function *SNP generator*, which, by default, extracts sequences of a user-defined length centred on all SNPs subsampled by the function *SNP Subsampler*. Here, for any probe, the base of SNP site will be replaced with one other than the two alleles provided in the input VCF file, to minimise potential bias towards the existing haplotypes. [38].

#### 2.2.4 Probe set assessment and validation

*eProbe* supplies several functions for assessing the final probe set. Function *SNP Assessor* allows for visibly profiling and generating statistics on the biophysical and biochemical features (see Supplementary Information), which form the fundamental feature of the synthesised capture baits. To assess if the probe set reflects population structure composed in the input VCF file, *SNP Assessor* can also generate hierarchical clustering heatmaps using pairwise hamming distance matrices on both the original input VCF file and the subset VCF file that only contains SNPs covered by the probe set, and then compute correlation coefficient and Manhattan distance between the two matrices (see Supplementary Information). *SNP Assessor* can also evaluate the effects of missing rate on the genome coverage of the captured data by simulating varying degrees of missing rate on probe-covered loci, and perform genome-wide Principal Component Analysis (PCA) to assess the robustness of genotyping with different levels of missing rate (see Supplementary Information).

### 2.3 Probe set for the FUNCGEN panel

For the FUNCGEN panel, eProbe provides the *Seq_generator* function, which generates probe sequences by tiling over the provided gene sequences based on the specified probe length and interval. This function accepts gene sequences in various forms (as described in the Input Data section). *eProbe* can also handle multiple haplotypes of a gene, by generating a probe for each different alleles (see Methods), either through user input or by estimating them using a VCF file containing the variants on target genes, thereby improving the sensitivity in detecting the genotypes for known genetic diversity.

## 3. Benchmarking experiment

We conducted a benchmarking study using samples from extant foxtail millets and wild foxtail millets (now collectively referred to as millets) to access the *eProbe* workflow, which also works as an instruction tutorial. A total of 50,000 probes were designed. After removing the duplicates 49,364 probes were synthesised as the capture bait set, including both POPGEN and FUNCGEN panels. For the POPGEN panel, 9.7 million bi-allelic SNPs distributed across the millet full genome were collected as the input dataset, with 32,357 probes finally retained to cover the same number of target SNPs (see Methods). An additional 17,643 probes were synthesised to cover 46 key genes (see Supplementary Information) associated to the millet domestication, as the FUNCGEN panel. Surface soils and leafs of millet plant specimens from three separate millet farms (Beijing, China) cultivating three different millet varieties [cultivar (*Setaria italica*, referred to as C hereafter), landrace (*Setaria italica*, referred to as L hereafter), and wild (*Setaria viridis*, referred to as W hereafter)] were shotgun and capture sequenced, yielding a total of 74,000,294 and 82,853,400 sequencing reads respectively, after data quality controls (see Supplementary Information). Taxonomic profiling was performed following the established pipeline for aeDNA (see Methods).

### 3.1 Remarkably increased sequencing efficiency

Most millet sequences (∼75%), whether from soils or plant tissues, originated from the conserved genomic regions in both *S. italica* and *S. viridis*. As a result, they were assigned to the ancestral node, i.e., the genus *Setaria* (see Supplementary Information). This supports the rationale for considering all sequences assigned to the *Setaria* genus as successfully enriched target sequences. Capture sequencing increased the average endogenous rate of millet by 455-fold compared to shotgun sequencing, ranging from 244-fold to 577-fold (Fig. 2a). In the shotgun dataset sequenced from soils, on average only 0.17% reads were assigned to *Setaria* genus or below, whereas in the capture dataset at least 50% total sequenced reads were assigned below *Setaria* (see Supplementary Information). Nearly all genome regions targeted by probes (hereafter referred to as target regions) were covered by the capture sequencing data generated from soil samples, with an average of over 20x coverage per gigabase of sequencing data. This marks a comparable average depth to the data captured from the plant tissues, and a significantly higher depth compared to the whole genome sequencing of plant tissues (Fig. 2b). The coverage of soil capture sequencing data also yielded uniform distributions across the target regions, with only occasional outliers of notably high depth on some regions (Fig. 2c). These all mark remarkable improvements on sequencing efficiency and genome coverage using the designed capture probe set compared to shotgun sequencing. Intriguingly, we found our capture probes yielded higher endogenous rate for the cultivar, presumptively due to the relatively higher millet DNA contents in the surface soil of the cultivar farmland, since the same higher rates were also observed in the shotgun sequencing data of the cultivar farmland (see Supplementary Information).

**Fig. 2.**
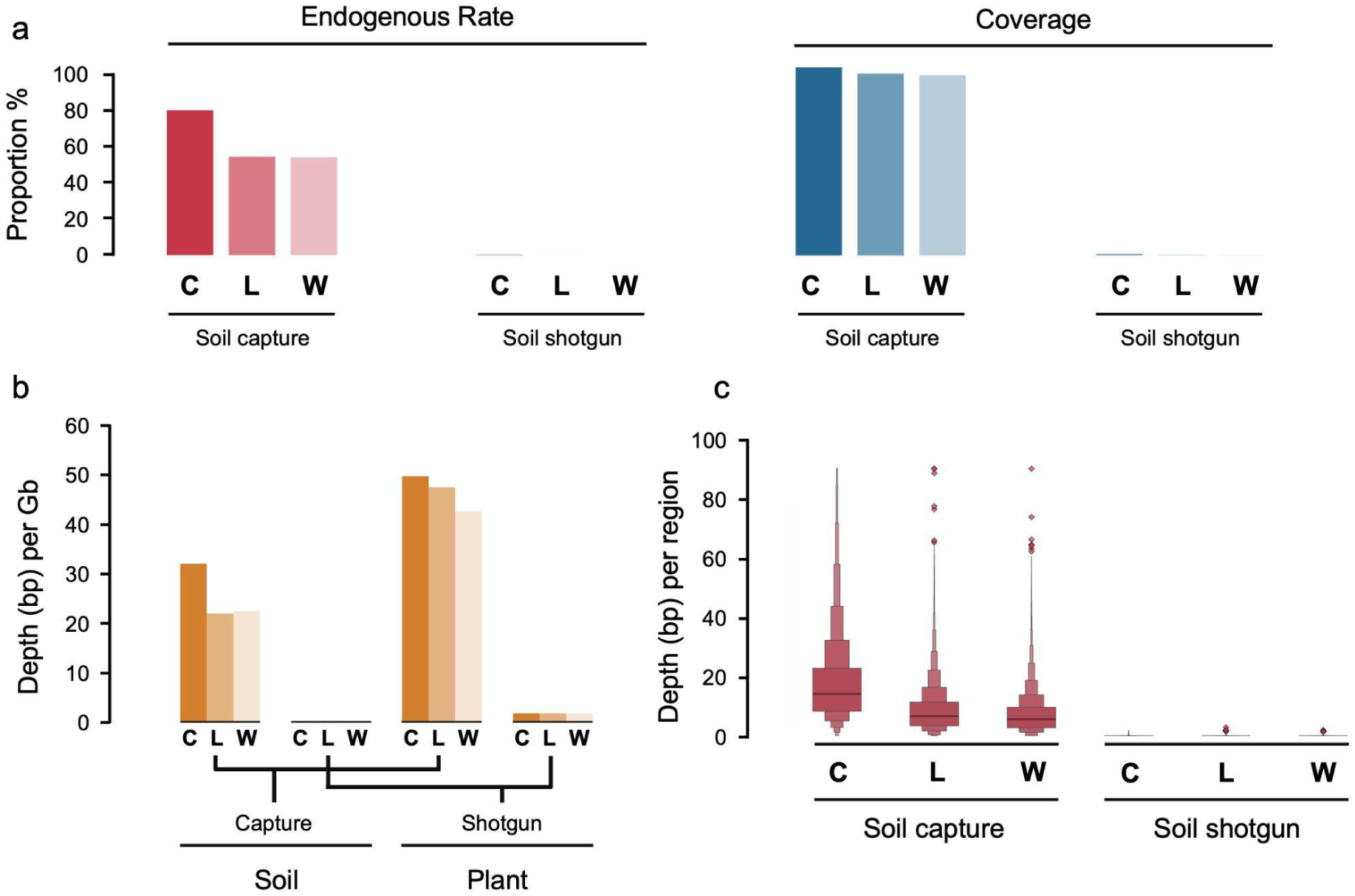
Benchmarked efficiency of the designed capture probes. (a) Millet endogenous rate and coverage rate on target regions (genome regions targeted by probes) for capture and shotgun sequenced data yielded from different millet farmland suffice. C stands for millet cultivar (*Setaria italica*), L for a millet landrace (*Setaria italica*), and W for wild millet (*Setaria viridis*). (b) Sequencing depth on target regions per gigabase sequencing data. Capture and shotgun data sequenced from both millet surface soil and plant specimens are shown. (c) Distribution of coverage rate on target regions. Width of the box indicates the number of target regions within each range of the coverage depth rates. The thick black line indicates the median of the distribution, while points indicate outliers.

### 3.2 Effects of probe filterings and pooling strategies

We compared the capture sequencing data targeted by the POPGEN panel (undergone all probe filterings) and the FUNCGEN panel (direct output from the input data therefore unfiltered), to testify the effects of probe filterings on capture efficiency. Two different sequencing library pooling strategies prior to the capture reaction were also compared, to assess the effects of pooling multiple libraries (indexed with different indexing primers) for capture reactions (see Methods).

We found significant improvements resulting from probe filterings (Fig. 3). Across the 6 soil capture libraries, the average number of reads normalised onto each capture bait for the filtered probes are ∼2.9 times higher than for the unfiltered probes, and the improvements are observed evenly across the target regions. The efficiency differences between the 2 panels are likely ascribed to the different levels of homogeneity for the probes against other taxa. The unfiltered probes in the FUNCGEN panel are mainly functional genes that are conserved among many taxa, and the captured reads would therefore be more likely assigned to other taxa than *Setaria* thus classified to higher ancestral nodes by the competitive mapping approach we applied (see Methods), and hence not identified as millet sequences anymore.

**Fig. 3.**
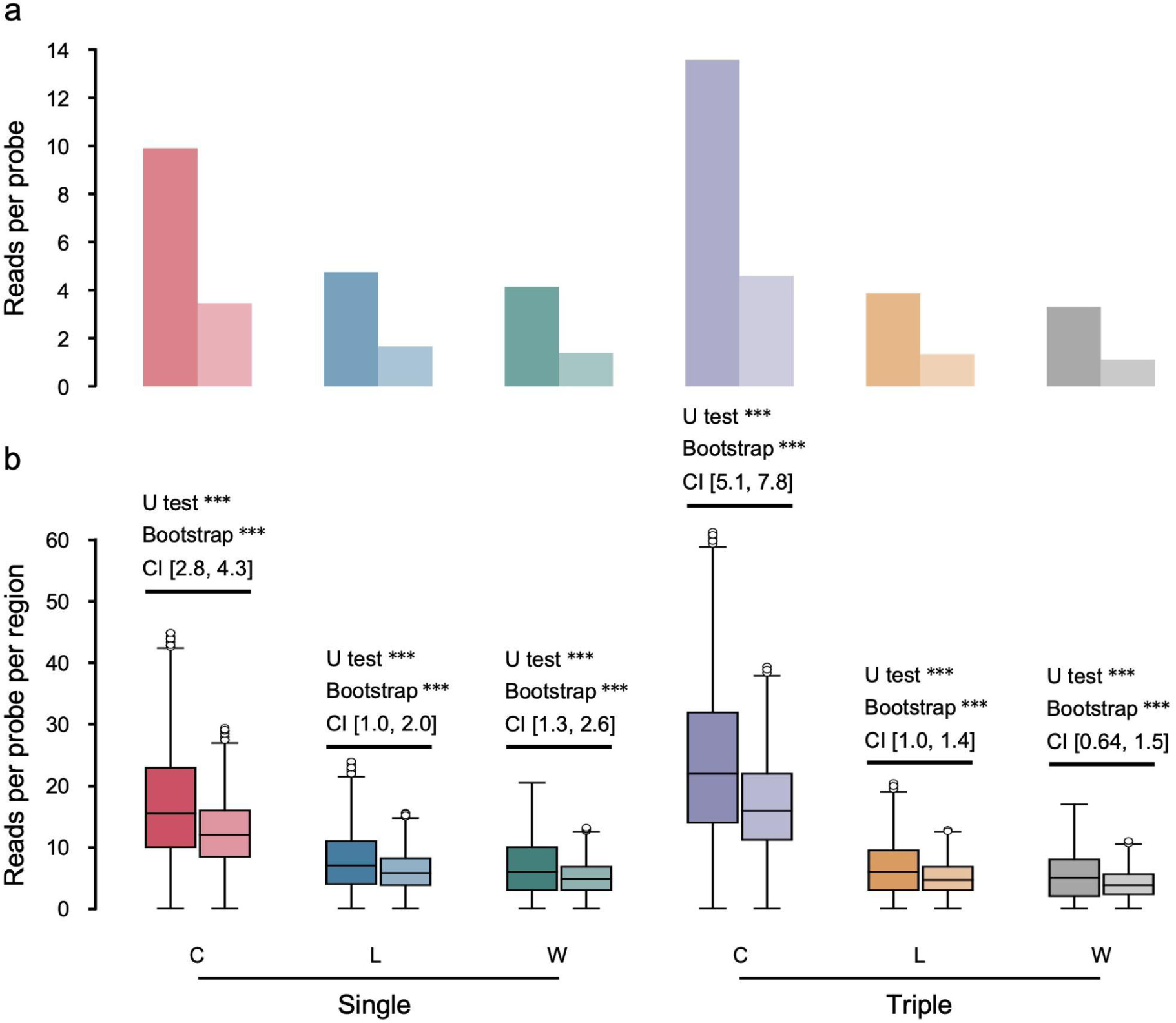
Effects of probe filterings and pooling strategies on capture efficiency. (a) The total number of reads overlapping target regions divided by the total number of probes, with the left bar in each pair denoting the filtered panel, and the right bar denoting the unfiltered panel. (b) The box plots show the number of reads overlapping each target region divided by the number of probes overlapping this region, with the left box in each pair denoting the distribution for the filtered panel, and the right box denoting the unfiltered panel. Single indicates a single library as input for the capture reaction, while Triple indicates 3 libraries indexed and pooled prior to the capture reaction. To enable comparisons between different libraries, the statistical results were normalised against the sequencing data size of each library. The Mann-Whitney U test and bootstrap resampling are depicted on each pair of boxes, indicating whether the capture efficiencies are statistically significant between the 2 panels.

When pooling different sequencing libraries for hybridization capture, we observed an overall increased yield of endogenous millet sequences compared to single library capture. However, we also found that most of the increases were contributed by the library derived from the foxtail millet cultivar sample, which has a higher millet DNA content (see Supplementary Information). This indicates potential bias can be introduced when pooling libraries with notable differences in the contents of the targeted sequences.

### 3.3 Effects of probes’ biophysical and biochemical features

We categorised the values of each biophysical or biochemical feature (averaged values of all probes on each target region) into consecutive intervals, and investigated the resulting coverage depth on each interval for each feature, to assess their effects on the capture efficiency (Fig. 4, See Methods).

**Fig. 4.**
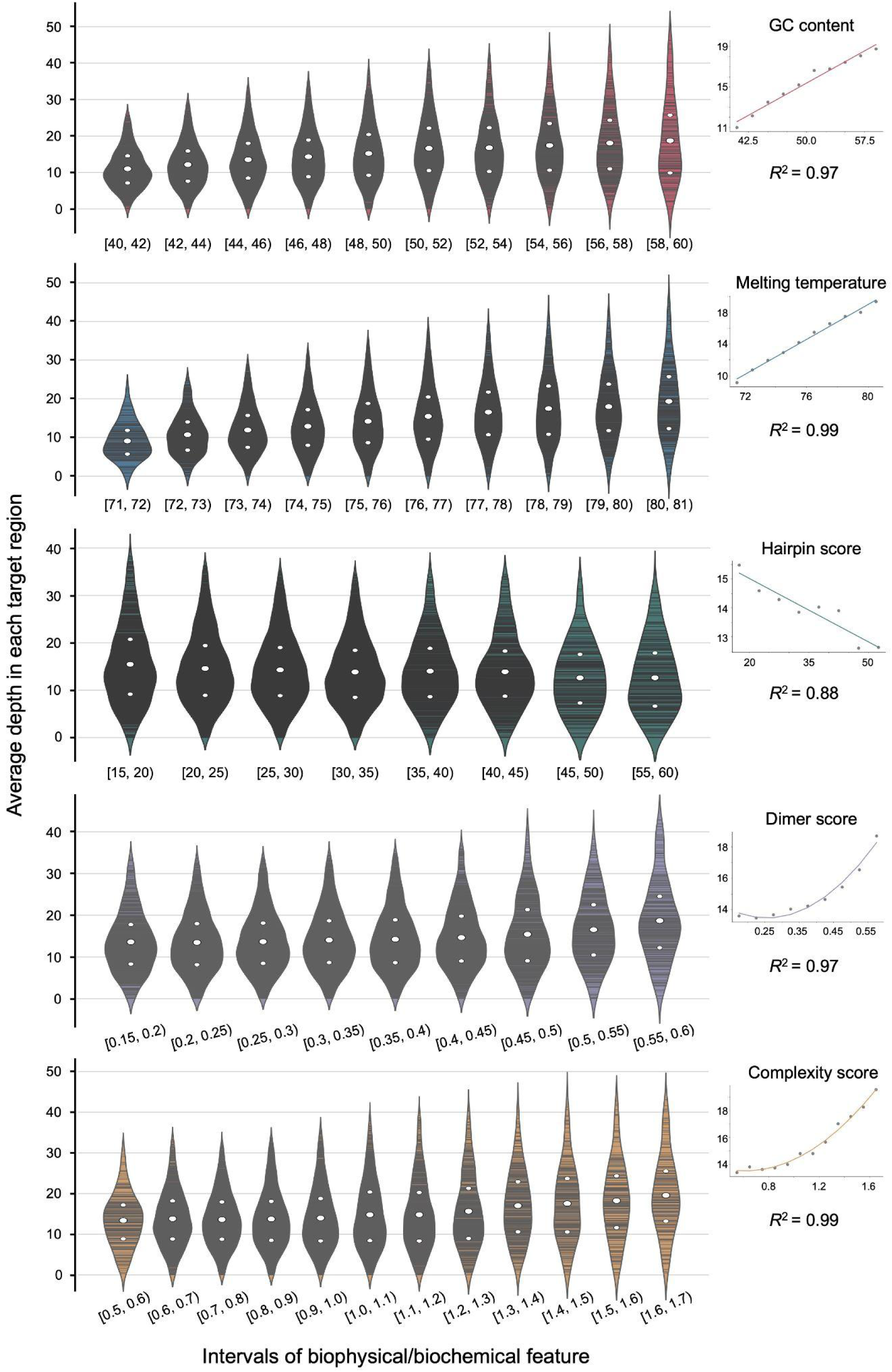
Effects of probes’ biophysical and biochemical features on capture efficiency. From top to bottom: the average depth of each target region against its GC content, melting temperature (Tm), hairpin score, dimer score, and probe sequence complexity (*S* score). Each violin box shows the distribution of average depth in an interval for the given feature. Each horizontal grey line within the violin box indicates the average depth of a target region. Correlations between the mean of average depth in an interval and the corresponding mean of biophysical or biochemical features are shown on the right for each feature.

Both the GC content and the Tm exhibit positive linear correlations to the captured coverage depth within a certain range, likely due to probes with higher GC content (typically also higher Tm) possess higher specificity and stability in hybridization binding [61]. However, similar to PCR primer design, extremely high GC content with excessive Tm would result in difficulties in hybridization binding and reduced tolerance in mismatches between probes and template eDNA sequences [62,63]. Therefore, moderate GC content and Tm that well balance the stability/specificity and the hybridization binding efficiency are recommended.

As expected, the hairpin score (marking the chances in forming within-probe hairpin structures) demonstrates a negative linear correlation to the coverage depth, indicating that the formation of within-probe hairpin structures could largely reduce the availability of probes and thus the capture efficiency. Both the dimer score (marking chances in forming dimmers among probes) and the probe sequence complexity (*S* score, higher value indicating lower complexity) show similar correlations to the captured coverage depth, that positive effects start to increase after passing certain thresholds. Notably, high dimer scores and low complexity could both indicate the presence of poly-X sequences and/or tandem repeats in the probe sequences, which are prone for nonspecific binding and thus increase the chance of cross binding of probes and template eDNA sequences from different genome regions. This suggests that within the filtered ranges (probes with outlier values for these 2 features had been filtered out before bait synthesis), high dimer scores and low sequence complexity seemingly do not decrease the capture efficiency.

Importantly, we note that these biophysical or biochemical features are not independent. The potential auto-correlations among different features, complicated working mechanisms, and their associations with other uninvestigated features as well as the factors involved in the hybridization reactions all require further studies for better understanding the patterns. Nevertheless, these results highlight the importance of the probe filterings that the *eProbe* workflow supplied.

### 3.4 Population genetic and functional gene analysis

To assess whether the data capture-sequenced from soil samples can reflect the millet population patterns, we performed population genetic analysis, including Identity-By-State (IBS) distance, PCA, and Admixture analysis, and compared the results between soil capture sequencing data and plant whole genome sequencing (WGS) data recovered from same millet farmlands (Fig. 5, See Methods). We also presented a preliminary result on a validated key domestication trait seed shattering, controlled by gene *shattering1* (*sh1* [64]), which was targeted by our FUNCGEN panel, to evaluate whether the probe set can effectively capture the polymorphisms of functional genes.

**Fig. 5.**
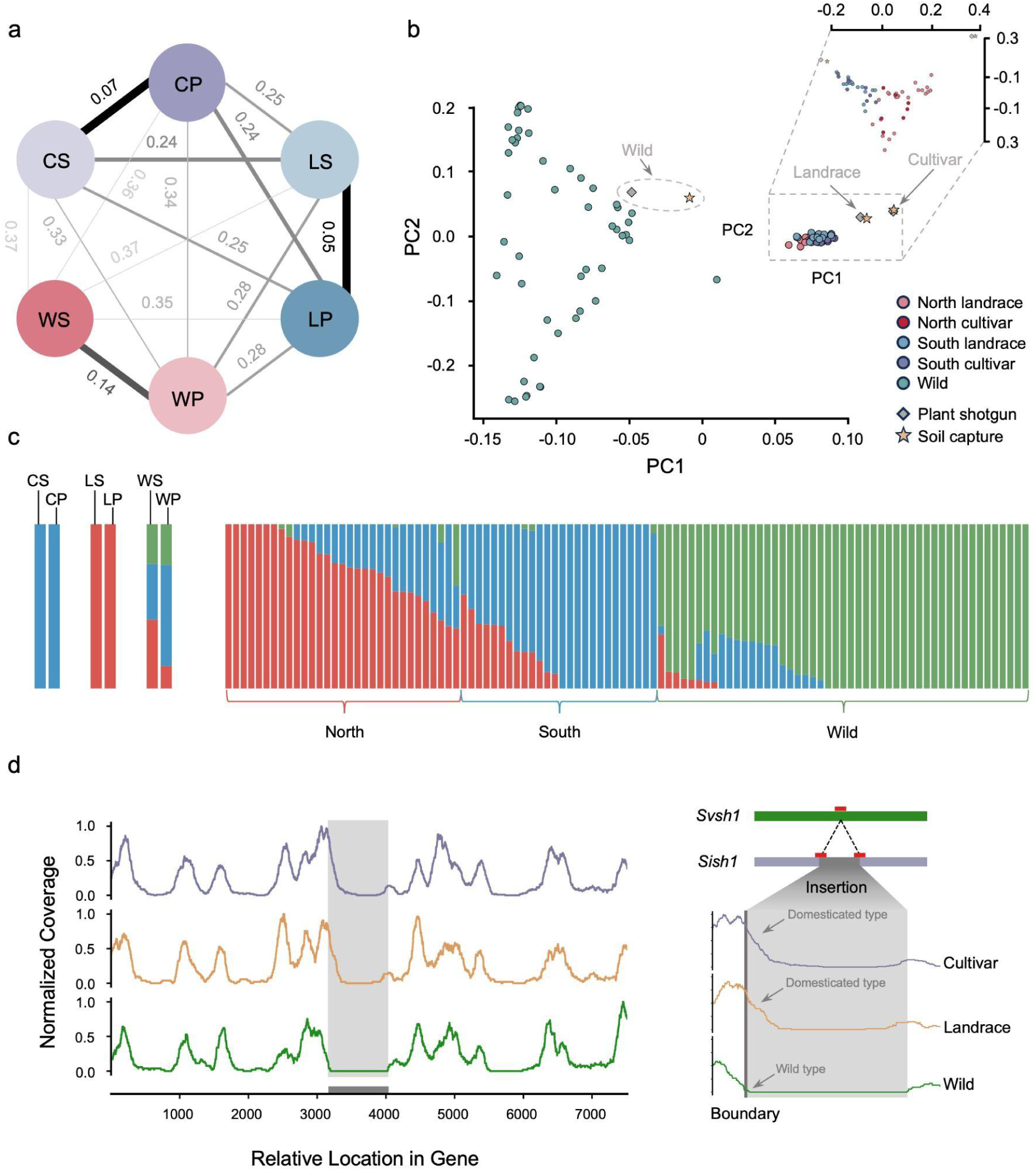
Population genetic and functional gene analysis on the captured millet DNA. The first letter in a sample ID denotes the millet variants, with C for cultivar, L for landrace, and W for wild, while the second letter denotes the data source, with S for soil capture sequencing and P for plant whole genome shotgun sequencing. (a) Pairwise Identity-By-State distances for all the 6 tested samples. Number on each line indicates the distance (1-IBS) between that sample pair. Line thickness is proportional to the distance. (b) Principal component analysis (PCA) including published wild and domesticated foxtail millet, and our soil and plant samples. (c) Admixture analysis showing the ancestral components of each sample. (d) Coverage landscape for the *sh1* gene. The left figure shows the coverage landscape of the complete *sh1* gene, with the bottom grey bar indicating the transposon insertion. The right figure further zooms into the insertion region, where the insertion is present in the *Sish1* haplotype (domesticated millet) but absent from the *Svsh1* haplotype (wild millet). The red bars indicate where the probes were placed to capture this variant.

Capture sequencing data recovered from the soils exhibit very small IBS distances to the corresponding cultivated plants (Fig. 5a), indicating that soil capture sequencing effectively captures the genetic patterns. The relatively bigger value of distance between the soil and plant samples for the wild millet is presumably due to this farmland being used for cultivating millet cultivars in previous years, the soil thus preserving a mixed DNA pool of both wild millet and domesticated millet cultivars. The distances between soil capture sequencing data for cultivar and landrace, and between plant WGS data for cultivar and landrace are almost identical, further indicating that our capture bait set well captures the genetic patterns of millet.

PCA and Admixture analysis further confirm the effectiveness of our capture bait set on resolving the population genetic patterns. The soil samples from the cultivar and the landrace farmlands largely overlapped with their corresponding plant specimens on the PCA (Fig. 5b). While the wild millet plant specimen was clustered together with other published wild millet genomes, its soil capture sequencing was placed in between the wild and domesticated panels, confirming that this sample contains mixed DNA of wild and cultivar millets. Same patterns were also observed in the Admixture analysis (Fig. 5c).

It is noteworthy that, due to the cultivar and landrace variants tested here are not regular millet variants represented by the reference genome panel, which was used for both probe designing and population genetic analysis, the recovered genomes from these variants did not cluster together with other genomes of domesticated millet. This, however, highlights the capability of our designed probe set for capturing unsampled genetic patterns of the target taxa, and the flexibility of *eProbe* workflow in avoiding potential bias introduced by the skewed reference genome database.

The *sh1* gene targeted by our FUNCGEN panel is a key gene associated with the millet domestication. This gene experienced a transposon insertion that led to the inhibition of natural seed shattering [64], and is therefore crucial in distinguishing between the domesticated and wild genotypes of millet. Our probe set effectively detected presences of the transposon insertion from the soil capture sequencing dataset of millet cultivar and landrace farmlands, while the insertion is absent from the data capture-sequenced from the wild millet farmland (Fig. 5d). This demonstrates the capability of our probe set in potentially detecting the structure variations for key functional genes, as well as distinguishing between domesticated and wild millet.

## 4. Conclusions

Following the recent dramatic growth of the field of ancient environmental DNA, it is clear that hybridization capture is central for facilitating genome-scale analyses for highly mixed and degraded DNA recovered from ancient environmental samples [20,38], which can provide insights into adaptive and demographic genetic processes beyond the constraints of sampling individual fossil remains. With the increasing number of applications of eDNA capture, a readily deployable and comprehensive program toolkit is required to efficiently and scalably design and access capture probes for target organisms. The *eProbe* package has been developed to meet this need, by providing a complete workflow to flexibly and optimally generate, filter, select and assess capture probes. The *eProbe* workflow facilitates taxa-specific population and evolutionary genetics analysis from environmental samples.

We demonstrated proof-of-concept by designing and synthesising a capture bait set for millet, and benchmarked its performance on surface soils sampled from farmlands cultivating various varieties of wild and domesticated millets. From our sequencing results, we show that the eProbe capture baits increased the sequencing efficiency by >450 fold. The application of probe filtering steps incorporated in the *eProbe* workflow significantly improved the efficiency of the capture baits. The capture-sequencing data recovered from soils yielded sufficient genome coverage depth on the millet genome, enabling detailed and accurate population genetic analysis. Probes designed targeting functional genes effectively captured the different genotypes of wild and domesticated millet from soil samples.

The *eProbe* package and workflow removes a major obstacle in the field of aeDNA, by supplying a dedicated tool for optimised *in silico* probe design and validation. This enables researchers from a wide range of backgrounds to utilise hybridization capture-based aeDNA analysis to their species of interest. Ultimately, this will increase accessibility to genome-scale data from aeDNA, facilitating research options in the evolutionary and demographical genetics using ancient environmental samples, and catalysing aeDNA into its genome era.

## 5. Methods

### 5.1 The *eProbe* workflow

#### 5.1.1 SNPs preprocessing

Function *SNP_preprocessor* employs the pysam package (v0.22, https://github.com/pysam-developers/pysam) to retrieve essential information from the input VCF file as the SNPs metadata, which is written as a SNPs tab-separated table (the SNPs TSV file) for subsequent processing. BEDtools v2.26.0 [65] is utilised to selectively retain or exclude SNPs delineated in the BED file (if supplied). SNP clusters are detected by counting the number of other SNPs around each SNP within a user-defined search length.

#### 5.1.2 SNPs filtering

Function *SNP_filter* performs all probe filterings. It first extracts sequences of a given length centred on each SNP on the reference genome as the raw probe candidates (as FASTA format), with storing SNP information in each probe sequence header. All probe candidates are then subjected to filterings, by modifying the SNPs TSV file to subset the target SNPs after each filtering. By default one probe will be generated targeting one filtered SNP.

*SNP_filter* incorporates kraken2 to match the probe candidates against the genome *k*-mer databases for background noise filterings. The filtering intensity is based on the minimum non-overlapping *k*-mer groups shared between each query to the supplied *k*-mer databases.

To filter inaccessible regions, *SNP_filter* incorporates bowtie2’s local alignment mode [66] to align probe candidates against the customised input genomes (normally the genomes used for SNPs calling to generate the input VCF file). *eProbe* provides two filtering modes: 1) strict mode: probe sequences must uniquely align to each genome (no secondary alignments) with a mapping quality ≥30, edit distance ≤3, and no gaps, and 2) moderate mode: probe sequences may have secondary alignments, but the secondary alignment score must not exceed 80% of the primary alignment score, with mapping quality ≥30, edit distance ≤3, and allowing gaps. *eProbe* also accepts customizations on alignment parameters.

A lowest common ancestor (LCA) clustering approach is used for filtering closely related taxa. A reference database compromising the genomes of relevant taxa is required, against which the probe candidates are aligned using bowtie2’s end-to-end multiple alignment mode. *SNP_filter* then calls *ngs*LCA [67] for LCA clustering for each probe candidate, and only the candidates assigned to user-defined specific ancestral nodes and their descendants are retained.

*SNP_filter* applies the functions from the Biopython [68] to calculate the GC content and melting temperature for each probe candidate. Estimation of melting temperature is based on the nearest-neighbour thermodynamic model [69]. Several tables containing thermodynamic nearest-neighbour values for DNA/DNA and DNA/RNA hybridizations are implemented, with the table for DNA/RNA hybridization [70] used as default. To assess sequence complexity, *eProbe* calculates the *S* score for each probe candidate [71]. By default, *eProbe* filters out sequences with an *S* score greater than 2, as these typically contain tandem repeats and poly-N structures.

To assess the likelihood of forming hairpin structures within a probe candidate, *SNP_filter* assesses the reverse complement degree within the sequence of each candidate. This is achieved by calculating alignment scores between each probe and its reverse complementary sequence using the PairwiseAligner module of Biopython running the local aligning mode and "blastn" scoring mode. The best alignment for each probe candidate were extracted, and hairpin score *H*(*a*) is estimated by formula:

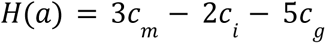

where *c_m_*, *c_i_*, and *c_g_*denote the base number of matches, mismatches, and gaps in the alignment, respectively. By default, *eProbe* filters out probes with a hairpin score greater than 60, which indicates that the probe contains two segments of a 10-bp sequence that can perfectly hybridise.

*SNP_filter* estimates the likelihood of probe dimer formation using a *k*-mer based approach. Probe candidates are chopped into *k*-mers meanwhile counting for the frequency of each unique *k*-mer to form a *k*-mer database. Each probe candidate is then reverse complemented to traverse all possible *k*-mer its sequence contains. A dimer score *D*(*a*) for probe *a* is defined by formula:

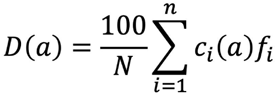

where *n* is the number of all possible *k*-mer that can be extracted from probe *a*; *c*_i_(*a*) is how many times a given *k*-mer *i* can be found in probe *a*; *f*_i_ is the frequency of the given *k*-mer *i* in the *k*-mer database; *N* is the number of total input probe candidates. By default, *eProbe* removes probe candidates with dimer scores in the top 15th percentile of all dimer scores.

#### 5.1.3 Assessing probe set

To examine the biophysical and biochemical features, *SNP_assessor* extracts probe sequences from the subsampled SNPs to calculate and summarise user-specified features, and then uses the Seaborn [72] to plot the distribution graphs for each feature.

Pairwise IBS distance matrices for both the original input genome-wide SNPs and the probe-covered SNPs are computed using PLINK v1.9 [73]. The Seaborn is then applied to generate heatmaps for distance matrices, with the fastcluster [74] employed for hierarchical clustering. The correlation coefficients (Pearson or Spearman) and the Manhattan distances between the matrices are calculated using functions from the Scipy [75].

The resilience of the probe set to missing data rates is evaluated by simulating varying degrees of missingness randomly distributed across the probe-covered SNPs. *SNP_assessor* calls PLINK to perform PCA on the SNPs before and after simulating the missing rates.

#### 5.1.4 Probe generation for FUNCGEN panel

Function *Seq_generator* employs a sliding window approach for tiling probes across the sequences based on the specified window size and step for the supplied functional genes. To make probes targeting different alleles, *Seq_generator* calls ShapeIt [76] and PyBedTools [77] to infer genotypes for different genes specified by the user (supplied as a BED file), based on a VCF file that contains the variants on the supplied genes. *Seq_generator* will then extract sequences from the reference genome and replace the variants with different alleles to generate probes for each of the alleles. *Seq_generator* can also handle user-specified FASTA files that contain haplotype sequences. For both eProbe-generated or user-input haplotype sequences, *Seq_generator* aligns (genome-alignment) them using CLUSTAL v2.1 [78], and then detects the variations when sliding across the aligned sequences. Multiple probes will be generated on sites where mutations are found, with each probe targeting a different mutation, while a single probe will be generated in regions without mutations.

### 5.2 Benchmarking experiment

#### 5.2.1 Millet probe design

The detailed approaches covering all steps of the full workflow described here can be found in the *eProbe* GitGub repository and the Supplementary Information.

A millet reference genome database was assembled covering its modern genetic diversity (details in Supplementary Information). For the POPGEN panel, the minimum and maximum values for SNP count covered by each probe were set to 1 and 5, respectively for filtering SNP clusters. The "--minimum-hit-groups" parameter was set to 2 for background noise filtering against a dataset composed of all microbial genomes (bacteria, fungi, archaea, and protozoa) from NCBI RefSeq database (Jan. 2023). The "moderate" mode was applied for filtering inaccessible regions against genome ME034V. We assembled a grass family genome database, including 66 genomes under the Poaceae family (details in Supplementary Information), for closely related taxa filtering with minimum and maximum edit distances as 0 and 2, respectively. From each probe generation sliding window, the probe with the optimal GC content and melting temperature was selected and further down-sampled to a total of 32,357 probes (100 bases long) as the final POPGEN panel.

For the FUNCGEN panel, 46 key genes related to millet domestication (see Supplementary Information) were targeted. BLASTN [79] was used to identify exons associated with domestication in millet, using homologous sequences of known domestication-related genes in rice as the references. Gene *SiSh1* [64] and *SiLes1* [80] were further validated for targeting both the wild and domesticated genotypes. The FUNCGEN panel was generated using all these exon sequences, yielding a total of 17,643 probes, which were merged with the POPGEN panel to form a final probe set of 50,000 probes.

#### 5.2.2 Sample collection, hybridization capture and DNA sequencing

Both plant specimens and surface soil samples were collected from 3 millet farmlands of the Millet Genetic Resources Laboratory Experimental Farmlands, Center for Crop Germplasm Resources, Institute of Crop Sciences, Chinese Academy of Agricultural Sciences, Beijin, China, in August 2023. The 3 farmlands each cultivated an improved cultivar of *Setaria italica* (Yugu18), a landrace of *Setaria italica* (SSR57) originated from Bulgaria, and a wild accession of *Setaria viridis* (A10) originated from the United States.

Fresh plant leafs were ground in liquid nitrogen. Plant DNA was then extracted using the Plant Genome DNA Rapid Extraction Kit (Sangon Biotech) following the manufacturer’s instruction. Soil DNA was extracted using the DNA-easy PowerSoil Kit (Qiagen) following the manufacturer’s instructions. All DNA extracts were fragmented, end repaired, linked with sequencing adapters, purified and went through prePCR using Enzyme Plus Library Prep Kit (iGeneTech, Beijing, China).

Out of the 50,000 probes initially designed, after removing the duplicated sequences (found in the unfiltered FUNCGEN panel), a total of 49,364 RNA baits were successfully synthesised by iGeneTech Bioscience Co., Ltd. (https://design.igenetech.com/). We performed single-library capture and triple-library capture (by pooling the three libraries used in single-library capture into one before hybridization) using 750 ng of sequencing library per sample, each with a unique index. We conducted the capture-enrichment experiments following manufacturer’s instructions (iGeneTech, Beijing, China). First, the pre-captured library solution was dried in a SpeedVac system (Thermo Fisher SPD120). For hybridization, 30 μL Hybridization Master Mix, which includes 13 μL TargetSeq One® Hyb Buffer v2, 5 μL Hyb Human Block, 2 μL TargetSeq® Blocking Oligo, 5 μL RNase Block, 3 μL Nuclease-Free Water and 2 μL TargetSeq® RNA Probes, was added into the dried pre-capture library and incubated in 80℃ for 5 minutes and 50℃ overnight (16 h) on a thermal cycler. Before capture, TargetSeq® Cap Beads, Wash Buffer 1 and TargetSeq One® Wash Buffer 2 v2 were mixed separately and centrifuged. 50 μL of TargetSeq® Cap Beads was washed for three times with Binding Buffer and then mixed with a 180 µL of Binding Buffer. For target capture of libraries, the 180 μL of TargetSeq® Cap Beads solution was transferred to the hybridization reaction mixture, and pipeted to mix well on a thermal cycler at 50℃. After that, the target captured library mixture (which includes the TargetSeq® Cap Beads solution and hybridization reaction mixture) was incubated on a mixer, and then the target captured library beads was separated from the supernatant with a magnetic stand. To remove the uncaptured DNA, the target captured library beads were washed with Wash Buffer 1 and pre-heated (50℃) TargetSeq One® Wash Buffer 2 v2. for 3 times. Afterwards, 80% ethanol was used to wash the beads and Nuclease-Free Water was used for the target captured library DNA dissolution. To meet the DNA sequencing requirements, post-PCR amplification was done for the target captured library with the Post PCR Primer and Post PCR Master Mix under the suggestion program. After purification and quantification, the target captured libraries were subsequently sequenced on the Illumina novaseq6000 platform running 150 paired-end mode.

#### 5.2.3 Data analysis

A total of 160,253,688 raw reads were generated and adapters were removed using AdapterRemoval v2 [81]. Fastp v0.23.2 [82] were used to further remove adapter residues (--detect_adapter_for_pe), deduplicate (--dedup), discard low complexity (--complexity_threshold 30) and ultra short sequences (-l 30). After data quality controls, 156,853,694 reads were retained for the 15 sequenced libraries (see Supplementary Information). Quality-controlled reads were then aligned to the grass family database mentioned in section 5.2.1, using bowtie2’s end-to-end mode with the "-k" parameter setting to 1000. The resulting BAM files were sorted using Samtools v1.17 [83], and taxa profiled using *ngs*LCA with the minimum and maximum edit distances setting to 0 and 2. Only reads classified belong to or below the *Setaria* genus were considered as millet reads and used for subsequent analysis.

The millet reads were re-mapped against the foxtail millet reference genome of cultivar *xiaomi* v1 [84] using BWA v0.7.17 [85] in "mem" mode. The resulting SAM files were filtered for reads with a mapping quality ≥30 using Samtools and deduplicated using Picard (http://broadinstitute.github.io/picard/). The "stats" function of SAMtools was used to count the number of reads mapped to the reference genome and calculate the endogenous rate (number of reads uniquely mapped on the reference genome divided by number of total quality-controlled reads) on the generated BAM file. The "intersect" module of BEDtools was employed to tally the number of reads mapped on the target regions, and the "coverage" module was used to calculate the overall coverage, average depth, and depth of each target region.

To assess the effects of probe filterings, efficiencies of probes generated from the POPGEN panel and from the FUNCGEN panel were compared, with overlaps between the 2 panels excluded. The total number of millet reads was divided by the number of probes for each panel as an overall assessment. The number of reads assigned onto each target region was retrieved and divided by the number of probes targeting that region, as the normalised efficiency per probe per target region, which were then tested using the Mann-Whitney U test and bootstrap resampling (bootstrap number 1000) across target regions. Here the same number of regions were randomly sampled from both panels for each run, and the median value of the unfiltered panel was subtracted from the median value of the filtered panel, and the *p*-value was calculated by counting the number of times that the difference was greater than 0.

To investigate the effects of biophysical and biochemical features, we first calculated the average depth of genome coverage for each target region using the "depth" module of Samtools. The coverage depth was then normalised for each sequencing library against its sequencing depth. The distributions of coverage depth were thereafter examined against the calculated values for the biophysical and biochemical features for each target region. Values for each feature were subsequently categorised into predefined intervals, requiring each interval containing a sufficient number of values. For GC content, melting temperature, and hairpin score, we employed linear regression to model the relationship between the feature and the coverage depth. For the sequence complexity and dimer score, which demonstrate non-linear correlation, polynomial regression was used for fitting the relationship.

SNPs calling was performed on the single-library captured data and their corresponding plant tissue sequencing data, for both soil samples and plant specimens, using BCFtools v1.3.1 [86]. The VCF file was filtered using VCFtools v 0.1.17 [87] to retain only biallelic variants (--min-alleles 2 --max-alleles 2) with a minor allele frequency ≥ 0.05 (--maf 0.05). IBS distances between each pair of the six samples were computed based on the filtered VCF using PLINK. We then merged the filtered VCF and that generated with resequencing data (details in Supplementary Information) and subsampled the SNPs located within the target regions from the millet reference genome database built for probe design. PCA was then performed on the merged VCF file using PLINK. ADMIXTURE [88] analysis was performed to infer the genetic admixture from different ancestral components.

## Acknowledgement

The authors thank Richard Durbin for the insightful discussions into this work. This work was supported by the Open Research Fund of TPESER (No. TPESER202202), the CAS Youth Interdisciplinary Team Fund, the NSFC BSCTPES Project (No. 41988101), the National Natural Science Foundation of China (No. 42101150), the Carlsberg Foundation (CF18-0024), and the Danish National Research Foundation (DNRF174). Data analysis was supported by the National Key Scientific and Technological Infrastructure project “Earth System Numerical Simulation Facility” (EarthLab, 2023-EL-ZD-000111), National Supercomputer Center in Wuxi utilising the computational resources of the Sunway TaihuLight Supercomputer, and SMU’s Center for Research Computing.

## Author contribution

Conceptualization: Y.W., H.W., Z.H.; Package development: Z.H., Y.W.; Benchmarking materials: Z.G., S.T., X.D.; DNA preparation: Z.G.; Data analysis: Z.H., Z.G., Y.C., H.D., Y.W.; Writing: Z.H., Z.G., R.M., Y.W., H.W.; All authors reviewed and commented on the submitted manuscript.

## Data and code availability

The sequence data generated in this study are available at BioProject PRJNA1140417. The *eProbe* package is under a GPL-3 licence (Non-Commercial): https://github.com/YCWangLab/eProbe.

